# Rotationally Stable Dynamics Over Long Timescales Emerge in Neuronal Development

**DOI:** 10.1101/2024.12.31.630917

**Authors:** Dominic R.W. Burrows, Richard E. Rosch

## Abstract

Neuronal networks must balance the need for stable yet flexible dynamics. This is evident during brain development, when synaptic plasticity during critical windows enables adaptability to changing environments whilst ensuring the stability of population dynamics. The emergence of population dynamics that balance stability and flexibility during development is poorly understood. Here, we investigated developmental brain dynamics in larval zebrafish, using *in vivo* 2-photon imaging to record single-cell activity across major brain regions from 3-8 days post-fertilisation, a highly plastic period in which hunting behaviours are established. Our findings revealed region-specific trajectories in the development of such dynamic regimes: the telencephalon exhibited increased neuronal excitability and long-range correlations, alongside the emergence of scale invariant avalanche dynamics indicative of enhanced flexibility. Conversely, while other regions showed increased state transitions over development, the telencephalon demonstrated a surprising rise in state stability, characterized by slightly longer dwell times and drastically reduced angular velocity in state space. Remarkably, such rotationally stable dynamics persisted up to 5 seconds into the future, indicating the emergence of strong attractors supporting stability over long timescales. Notably, we observed that telencephalon dynamics were maintained near to but not at a phase transition, thus allowing for robust responses while remaining adaptable to novel inputs. Our results highlight regionally-specific trajectories in the relationship between flexibility and stability, illustrating how developing neuronal populations can self-organize to balance these competing demands.

**Significance Statement:** Brain networks must balance the flexibility to adapt to new stimuli with the need for stability. This trade-off is particularly important during periods of high plasticity in brain development. Our study investigates this balance by recording single-cell activity across the entire brain of developing larval zebrafish. We discovered that brain dynamics become increasingly diverse, characterized by both short and long bursts of activity, reflecting increased flexibility. Simultaneously, we observed the emergence of stable dynamics, linked to consistent activity patterns over time. Using a modelling approach, we showed that this stability was driven by the formation of stable attractors that shape the dynamic trajectories. These findings highlight how population mechanisms can shape the dynamic interplay between flexibility and stability in regional networks in the developing brain.

## Introduction

Neuronal networks exhibit a fundamental trade-off between stability and flexibility, a balance crucial for robust yet adaptable neuronal responses (Magnasco, 2022; Safron et al., 2021). This balancing act is evident through synaptic remodelling in development (Faust et al., 2021; Lohmann & Kessels, 2014; Tau & Peterson, 2010) – activity-dependent plasticity during critical windows ensures sensitivity to changing environments (Ganguly & Poo, 2013), while homeostatic plasticity regulates global synaptic strengths retaining dynamics within a stable range (Turrigiano & Nelson, 2004). This push and pull between synaptic strengthening and pruning facilitates the formation and dissolution of attractor states (Amari, 1972; Hopfield, 1982; Pereira & Brunel, 2018), stable patterns of neuronal activity towards which the systems’ dynamics flow (Khona & Fiete, 2022).

The emergence of large-scale *in vivo* population recordings has enabled inference of attractors in neuronal dynamics. In particular, studies have reported multiple types of stable dynamics, including bistable up-and-down states (Cossart et al., 2003; Jercog et al., 2017), multistable whole brain states (R. E. Rosch et al., 2024), and continuous attractors in head-direction (Chaudhuri et al., 2019; Rybakken et al., 2019) and grid cell circuits (Gardner et al., 2022). Such stable dynamics support robust neural responses in the face of noisy inputs, for example to facilitate pattern completion in memory retrieval (Rolls, 2013). Interestingly, another line of argument states that the brain exists in a weakly attracting regime, which resides between stability and chaos (Beggs & Plenz, 2003; Cocchi et al., 2017; Hesse & Gross, 2014; Ponce-Alvarez et al., 2018). In this way, residing at a phase transition allows the system to flexibly encode a diversity of inputs (Bertschinger & Natschläger, 2004; Kinouchi & Copelli, 2006; Shew et al., 2009), while also maximising the number of available states (Haldeman & Beggs, 2005). Taken together, these competing dynamical perspectives underscore an inherent tension between maintaining robust, stable responses and exhibiting adaptability to novel environmental stimuli (Hillar & Tran, 2018; Mosheiff & Burak, 2019). This tension is particularly evident in brain development, when regionally specific brain function emerges and places changing demands on brain dynamics.

During early brain development, synaptic remodelling can serve as both a stabilising force and a source of flexibility. For example, monocular deprivation during critical developmental periods results in the redistribution of synaptic strengths towards the open eye (Wiesel & Hubel, 1963). Here, developmentally-induced activity-dependent plasticity can widen the repertoire of available states according to changing inputs. Concurrently, genetically hard-wired programmes must guide network connectivity towards evolutionarily optimised stable states, as evidenced by robust developmental trajectories in the zebrafish tectum in the face of changing inputs (Niell & Smith, 2005). The interplay between synaptic scale stability and flexibility in neuronal circuits has been well studied. How stability and flexibility is balanced at the level of neuronal populations, and the emergent dynamics that enable this to take place over development is largely unknown. Theoretical and *in vitro* work has suggested the self-organisation of population dynamics to a phase transition across development (Tetzlaff et al., 2010; Yada et al., 2017), but this has yet to be tested in living systems. Given that maladaptive attractors give rise to impaired dynamics in neurodevelopmental disorders (D. R. W. Burrows et al., 2020, 2023), studying this flexibility-stability trade-off may be key to understanding the functional consequences of such developmental abnormalities.

In order to study the flexibility-stability trade-off in development we require access to i) single cell firing properties, to accurately assess emergent population dynamics and ii) entire population coverage, to minimise subsampling artefacts. With this in mind, the larval zebrafish is an ideal model system, due to its optical transparency and small size enabling recordings of single cell dynamics across most of the brain (Ahrens et al., 2013). Furthermore, larval zebrafish exhibit highly heterogeneous cell-type composition, connectivity and activity patterns across different brain regions (Kunst et al., 2019; Mu et al., 2019; Scott, 2009), whilst sharing neuroanatomical functionality with mammalian counterparts (Kalueff et al., 2014; Mathuru & Jesuthasan, 2013) making it ideal for studying emergent dynamics across brain regions. In this study we record single cell activity across four major brain regions: i) the *telencephalon*, including the pallium (analogous to cortex in mammals), subpallium and olfactory bulb, ii) the *diencephalon*, including thalamic regions and the habenula, iii) the *midbrain*, including the optic tectum and tegmentum, and iv) the *hindbrain*, including the cerebellum and medulla oblongata (Kenney et al., 2021). Here, we analyse the single cell and population dynamics that emerge over development across major brain regions to assess stability and flexibility, and make use of *in silico* network modelling to understand how this trade-off is achieved.

## Results

In this study we analysed spontaneous neuronal dynamics at single cell resolution throughout early development to examine the dynamic interplay between stability and flexibility in neuronal populations. To do this we performed *in vivo* 2-photon volumetric imaging of larval zebrafish, recording single cell activity across the telencephalon, diencephalon, midbrain and hindbrain, from 3-8 days-post-fertilisation (dpf) (See Methods) (**Fig.1A-C**). During this period, larval zebrafish establish independent visually-guided feeding behaviours, requiring specialised sensorimotor integration of visual input towards goal-directed motor output, whilst negotiating internal states such as hunger against environmental inputs.

**Figure 1.**
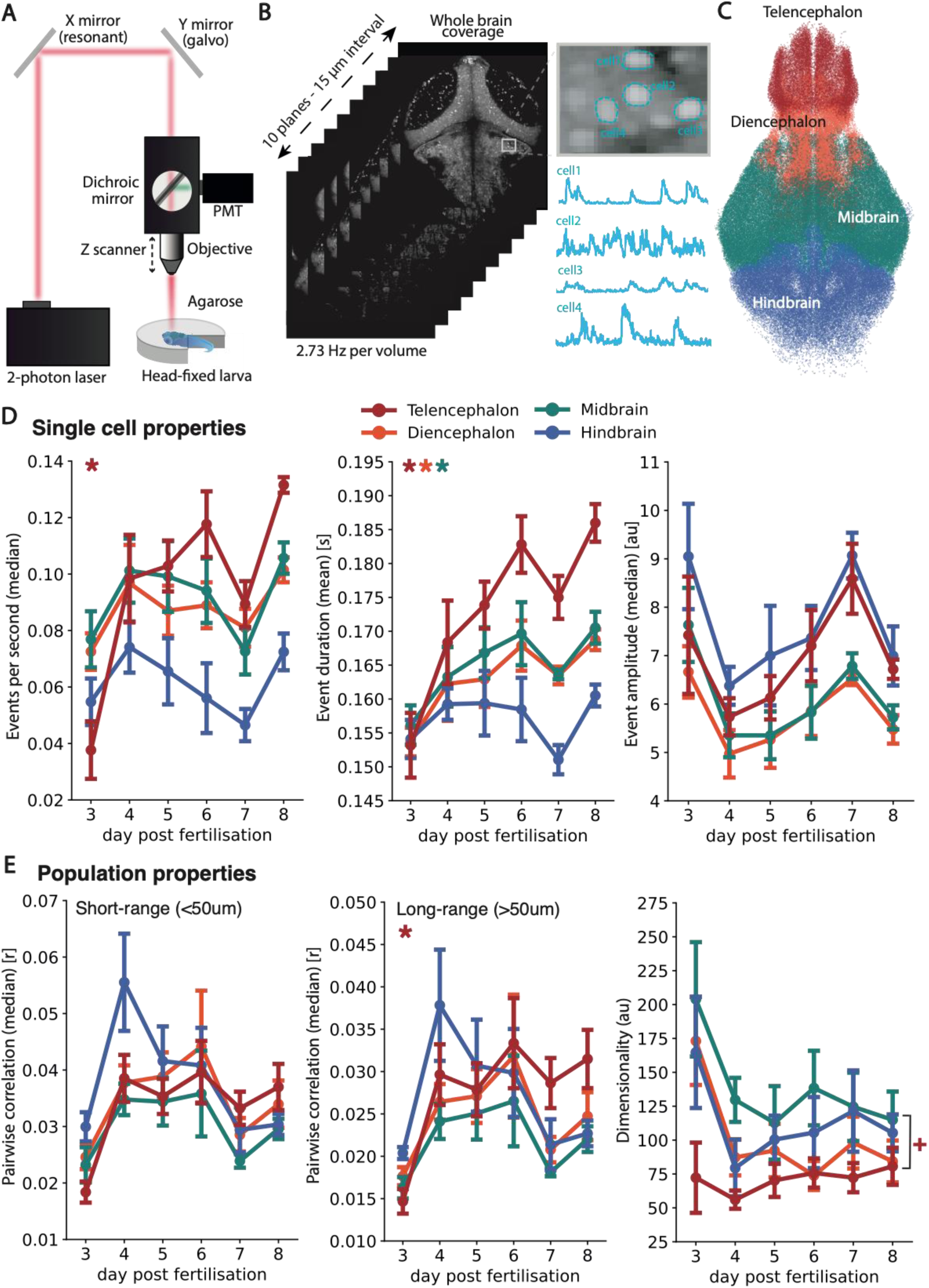
Neuronal firing properties across development. (A) In vivo 2-photon imaging setup. (B) 10 mean images from an example dataset, with enhanced region showing exemplar cells with nuclear fluorescence enabling nuclear segmentation, and fluorescence extraction below. (C) Single cell segmentation and regional labelling of neurons from the four major brain subdivisions, plotted from 14 animals across 6-8dpf. (D) Single cell properties quantified across major brain areas, showing the median events per second (left), mean event duration (middle) and median event amplitude (right). (E) Population properties quantified across major brain areas, showing the short-range median pairwise correlation (left), long-range median pairwise correlation (middle) and dimensionality (right). Data are significant with Bonferroni correction at α = 0.05 for Spearman’s test (*) or Mann Whitney U telencephalon vs other brain region pooled across ages (+). Error bars are standard error. Each datapoint is an animal, with means/medians taken across all cells for that animal.

### Neuronal Firing Properties Across Development

Population dynamics are a function of its underlying cellular activity. Therefore, to understand the constraints on population dynamics across development we first studied cellular firing properties. We first quantified the rate of calcium events, as a proxy for the spike rate of a given neuron. The event rate for each neuron was estimated as the number of timesteps classified as ‘on’ per second, by a Hidden Markov Model (HMM) designed for calcium fluorescence timeseries data (Diana et al., 2019). Here, we found that the telencephalon showed a significant, positive correlation between the median events per second and age (Spearman’s p=0.61, p < 0.001), increasing in event rate from 0.04±0.02 events/s at 3dpf, to 0.13±0.00 events/s at 8 dpf (**Fig.1D**). Other brain regions failed to show a linear correlation, but showed coherent non-linear dynamics - an increase in event rate from 3 to 4 dpf followed by a plateau, and then a further increase from 7 to 8 dpf. Next, we estimated the duration of calcium events, as a measure of the length of burst firing events within neurons. This was calculated as the mean length of each contiguous segment of HMM-labelled ‘on’ timesteps for a given neuron. Here, the telencephalon also exhibited a significant, near-monotonic increase in mean event duration with age (Spearman’s p=0.69, p < 0.0001), increasing in duration from 0.15±0.01 seconds at 3dpf to 0.19±0.00 seconds at 8 dpf (**Fig.1D**). Furthermore, the diencephalon and midbrain exhibited smaller, but significant positive correlations with age (Diencephalon: Spearman’s p=0.52, p < 0.01; Midbrain: Spearman’s p=0.52, p < 0.02). We also quantified the amplitude of calcium signals, as a measure of the intensity of burst firing events, which would give rise to high concentrations of intracellular calcium. Event amplitude was calculated as the median ΔF/F across active timesteps for a given neuron (see Methods). While no brain areas showed significant correlations with age, (telencephalon: Spearman’s p=0.21, p=0.27; diencephalon: Spearman’s p=-0.03, p=0.87; midbrain: Spearman’s p=-0.06,p=0.74, hindbrain: Spearman’s p=0.04, p=0.83), all regions exhibited remarkably similar dynamics – a reduction in amplitude from 3-4 dpf, followed by an increase from 4-7 dpf and a further reduction from 7-8 dpf (**Fig.1D**). Taken together, these results demonstrate that the telencephalon in particular exhibited markedly increased event rate and duration.

Another key statistical feature of population dynamics is the correlation between neurons in the network. To ascertain the correlational structure of neuronal networks throughout development, we quantified the pairwise correlation of neuronal activity across all neuron pairs. Specifically, we estimated the median Pearson’s correlation across all short-range (<50um) and long-range (>50um) connections, to disentangle the correlational structure of neighbouring and distant neuron pairs. While most regions showed no age effect across short and long-range connections, the telencephalon exhibited a significant increase in long-range correlations with age (Spearman’s p=0.47,p<0.01), from 0.01±0.00 3dpf to 0.03±0.01 at 8dpf (**Fig.1E**). This indicates increased coupling of activity across long spatial scales in the telencephalon across development. Interestingly, across both short and long-range correlations, the hindbrain exhibited a sharp increase from 3 dpf (short-range: 0.02±0.00; long-range: 0.01±0.00) to 4 dpf (short-range: 0.04±0.01; long-range: 0.03±0.01), followed by a step-wise decrease with age. In order to quantify the covariance across the entire population rather than across neuron pairs, we estimated the dimensionality of the dynamics (see Methods). Generally, high population covariance would lead to a lower dimensionality. While no brain regions exhibited significant linear age effects, the midbrain in particular exhibited a sharp decrease in dimensionality from 3-5dpf before plateauing, suggesting an increase in population covariance in early development. Interestingly, when pooling across ages, the telencephalon exhibited significantly reduced dimensionality compared to other brain regions (U=661, p<0.0001) (**Fig.1E**). This indicates higher population covariance in the telencephalon, compared to the rest of the brain.

Overall, we find that the telencephalon increases its single neuron excitability and long-range pairwise correlation throughout early development, while also exhibiting higher network covariance than other brain regions. The emergence of a more correlated, excitable state across development is suggestive of the organisation of the network towards a more synchronous and input-sensitive regime.

### Neuronal Avalanche Dynamics Across Development

Next, we set out to quantify the propagation of activity through the population across development. A useful approach for quantifying propagating dynamics is the neuronal avalanche framework (Beggs & Plenz, 2003), which describes how local bursts of activity spread through the network. Importantly, avalanche statistics can be used to examine the flexibility of population dynamics – avalanches that are scale invariant, spanning diverse time and length scales, support flexible encoding of inputs and a wide repertoire of accessible states (Haldeman & Beggs, 2005; Kinouchi & Copelli, 2006; Shew et al., 2009). Such a regime, known as criticality, occurs when the dynamics are organised to a phase transition (see Methods) (Bak et al., 1987; Perković et al., 1995; Sethna et al., 2001).

In order to assess the flexibility of population dynamics we calculated avalanche dynamics across development (see Methods). When we visualise empirical distributions for avalanche size and duration over development, we note that avalanches start off small in size and short in duration at 3 dpf, before evolving into much larger avalanches that span multiple orders of magnitude in size and time (**Fig.2A**). To estimate the slopes of the avalanche distributions, we fit power laws to our empirical distributions (see Methods). Here, we find that the size (τ) and duration (α) power law exponents significantly negatively correlate with age, across the telencephalon (τ: Spearman’s p=-0.66, p<0.0001; α: Spearman’s p=-0.64, p<0.001), diencephalon (τ: Spearman’s p=-0.53, p<0.01; α: Spearman’s p=-0.53, p<0.01) and midbrain (τ: Spearman’s p=-0.47, p<0.01; α: Spearman’s p=-0.58, p<0.001) (**Fig.2B**).

**Figure 2.**
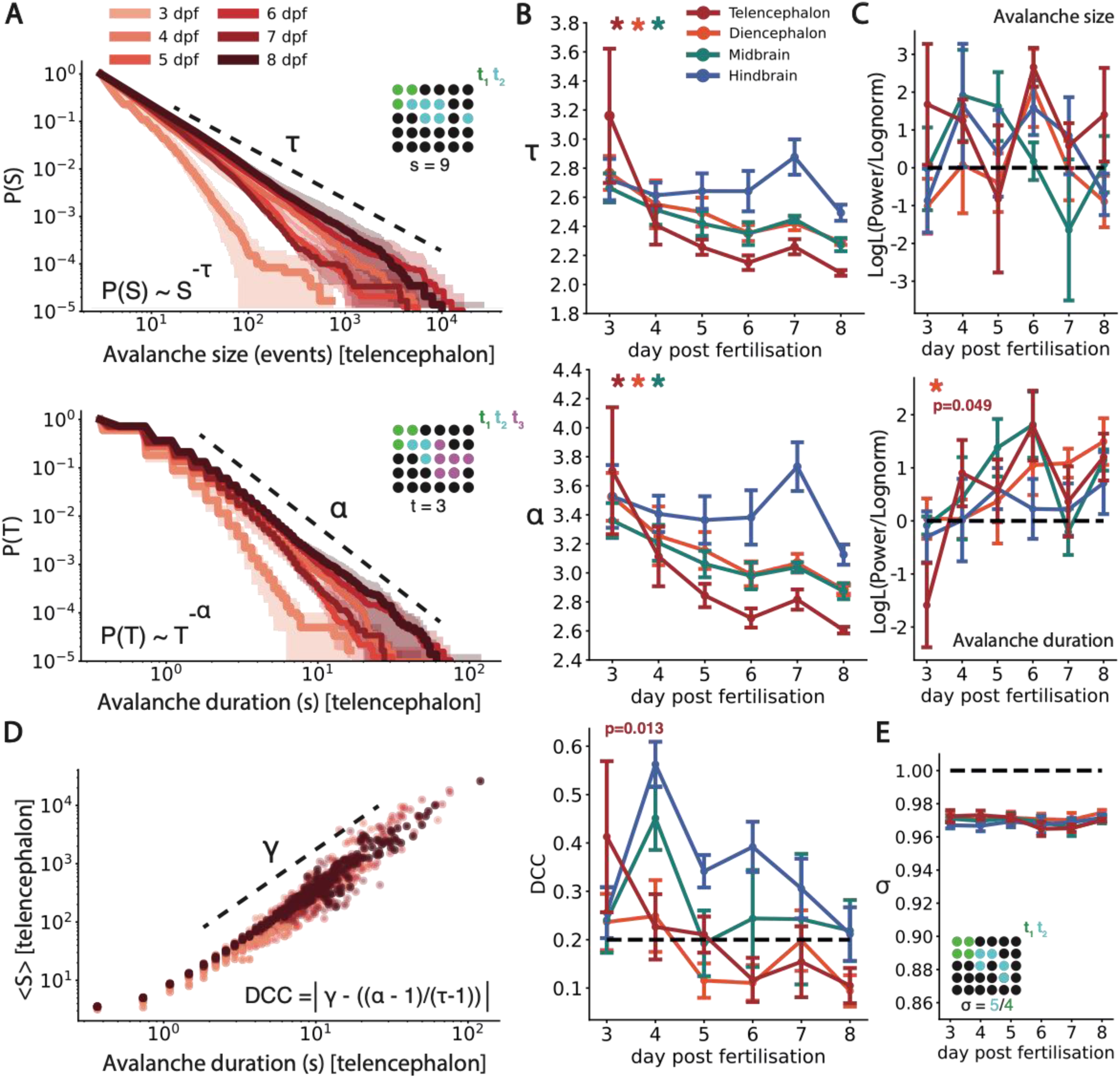
Critical avalanche dynamics across development. (A) Mean empirical distributions for avalanche size S (top) and duration T (bottom), shown for the telencephalon from 3-8 dpf. Avalanche schematic (Panel top right) demonstrating calculation of avalanche size (top) and duration (bottom) for single avalanche events. Black neurons are off. Coloured neurons represent active neurons at time point t_x_. (B) Mean avalanche exponents for size τ (top) and duration α (bottom) visualised across major brain subdivisions from 3-8 dpf. (C) Log likelihood ratios for power law vs log normal distributions for avalanche size (top) and duration (bottom), shown across major brain subdivisions from 3-8 dpf. Positive means indicate greater evidence for power law. (D) The scaling exponent γ captures the relationship between avalanche size and duration, here visualised as the mean size〈S〉for each duration. Notice how this follows a power law relationship at 8dpf, which breaks down at earlier ages. Each dot represents an〈S〉(T) relationship for a single fish. (E) DCC values are visualised across major brain subdivisions, from 3-8 dpf. (F) Branching ratio relationship σ visualised across major brain subdivisions, from 3-8 dpf. Branching process schematic showing approximate estimation of σ. σ is the number of descendants at a given time step (t_2_) divided by the number of ancestors (t_1_). Data are significant with Bonferroni correction at α = 0.05 for Spearman’s test (*). Error bars and shaded regions are standard error.

Overall, across development avalanche size and duration slopes decrease, indicating a higher propensity for larger and longer avalanches. Interestingly, the telencephalon shows the greatest change in avalanche slopes with age, from 3 dpf (τ: 3.16±0.81; α: 3.70±0.77) to 8 dpf (τ: 2.08±0.03; α: 2.61±0.04), mirroring observed neuronal excitability increases across development. Therefore, increased single neuron excitability leads to a concomitant expansion of propagating dynamics across development, particularly in the telencephalon. Notably, by 8dpf avalanches of all sizes from small to large occur with a greater frequency than at 3dpf (**Fig.2A**), indicating a greater diversity of propagating events across development.

To statistically test the presence of scale invariance in avalanche distributions, we performed log likelihood ratio testing (see Methods). For avalanche duration α, the diencephalon (Spearman’s p=0.50, p<0.01) exhibited a significant positive correlation with age (**Fig.2C**), while the telencephalon exhibited a moderate positive correlation which did not survive multiple comparisons correction (Spearman’s p=0.36, p=0.049). Nonetheless, at 3 dpf the telencephalon shows greater evidence for a lognormal distribution (LLR=-1.59±1.39), switching across development towards a power law by 8 dpf (LLR=1.20±0.77). This indicates that power laws in avalanche duration become more power law like across development in the diencephalon, which may occur to lesser extent in the telencephalon.

For avalanche size τ we find that the telencephalon exhibits greater evidence for power laws across all ages (LLR=1.08±1.03), except at 5dpf where the variance is particularly high (LLR=-0.82±3.48). Other brain regions exhibited more mixed evidence for power law distributions across all ages (diencephalon: LLR = -0.04±0.75; midbrain: LLR = 0.46±0.86; hindbrain: LLR = 0.47±0.84). However, we find no evidence of any age effects (telencephalon: Spearman’s p=0.03, p=0.87; diencephalon: Spearman’s p=0.08, p=0.68; midbrain: Spearman’s p=-0.08, p=0.68; hindbrain: Spearman’s p=0.08, p=0.67 (**Fig.2C**). Overall, observed avalanche expansion is most pronounced in the telencephalon, where avalanches become more diverse across development spanning both short and long time scales.

To further examine the scale invariance of avalanche dynamics, we characterised the relationship between the avalanche exponents τ and α. Another key property of scale invariant systems at criticality, is a scaling relationship between avalanche size *S* and duration *T* (Perković et al., 1995; Sethna et al., 2001), as

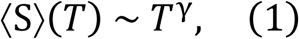

where *γ* captures a combination of avalanche exponents (see Methods). Put simply, the expected size of an avalanche should scale with its duration as a power law, with exponent *γ*. To assess this, we plotted the mean size of an avalanche〈S〉against its duration T. Here, we find that〈S〉(T) distributions follow an approximate scaling law, as evidenced by a log-linear shape which extends in length from 3 to 8 dpf (**Fig.2D**). Importantly, at criticality the relationship between the scaling exponent *γ* and avalanche exponents τ and α should follow:

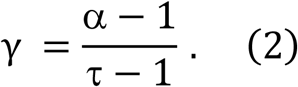

This exponent relation is a key property of scale invariant systems, and a strong indicator of flexibility in system dynamics (see Methods). Here we assess the presence of exponent relations using the deviation from criticality coefficient (DCC) (see Methods). Interestingly, we find that DCC values in the telencephalon exhibited strong negative correlations with age (Spearman’s p=-0.45, p=0.013), reducing from 0.41±0.27 at 3dpf, to 0.11±0.05 at 8dpf, marginally above the multiple comparisons threshold (**Fig.2E**). In contrast, no other brain regions showed clear age effects (diencephalon: Spearman’s p=-0.34, p=0.06; midbrain: Spearman’s p=0.19, p=0.30; hindbrain: Spearman’s p=-0.30, p=0.11). Therefore in the telencephalon, empirically observed scaling exponents γ seem to approach (α - 1)/(τ - 1) across development. This suggests the emergence of self-similar patterns in avalanche scaling relationships between *S* and *T*. Such scale invariant dynamics are suggestive of a wide repertoire of available system states, rendering the system more flexible and sensitive to inputs over development.

To build on these results, we model the dynamics as a branching process (Harris, 1964), which describes how an ancestor unit can generate descendant units, e.g. a neuron activating neighbouring neurons. Here, the phase transition separates a regime in which avalanches exponentially decay (subcritical), and one in which avalanches exponentially grow (supercritical). At the critical point, avalanches are scale invariant resulting in σ ∼1 (Harris, 1963). We use ‘MR. Estimator’, a toolbox which uses multiple-regression to estimate σ in a manner that is robust to subsampling (Spitzner et al., 2021). Interestingly, we find that all brain regions reside slightly below the critical point across development, (telencephalon: σ=0.97±0.00; diencephalon: σ=0.97±0.00; midbrain: σ=0.97±0.00; hindbrain: σ=0.97±0.00) (**Fig.2F**). In this slightly subcritical range, in the vicinity of a phase transition, the dynamics can be close to scale invariant while retaining slightly stable attractor states (Wilting et al., 2018).

### The Emergence of Rotational Stability Across Development

To further examine the flexibility-stability trade-off, we quantified the ability of the brain to flexibly transition across brain states. Here, we defined states as a set of state vectors (neuronal activity across the population at time *t*) that show non-random similarity to each other. To find such states, we clustered all state vectors for a given fish using affinity propagation and performed permutation testing on shuffled data.

Interestingly, while the diencephalon and midbrain showed a significant increase in the number of brain states across development (diencephalon: Spearman’s p=0.154 p<0.01; midbrain: Spearman’s p=0.49, p<0.01) (**Fig.3A**), the telencephalon showed no age effect (Spearman’s p=0.12, p=0.51). Surprisingly, the telencephalon exhibited an increase in the stability of its states, as shown by a moderate increase in the mean dwell time across all states (Spearman’s p=0.43, p=0.017), from 0.38±0.00 seconds at 3 dpf, to 0.40±0.00 seconds at 8 dpf (**Fig.3B**). Therefore, rather than an increase in flexibility of state transitions across development as expected in scale invariant dynamics, telencephalon dynamics show some evidence for increased stability of states through increased dwell time.

**Figure 3.**
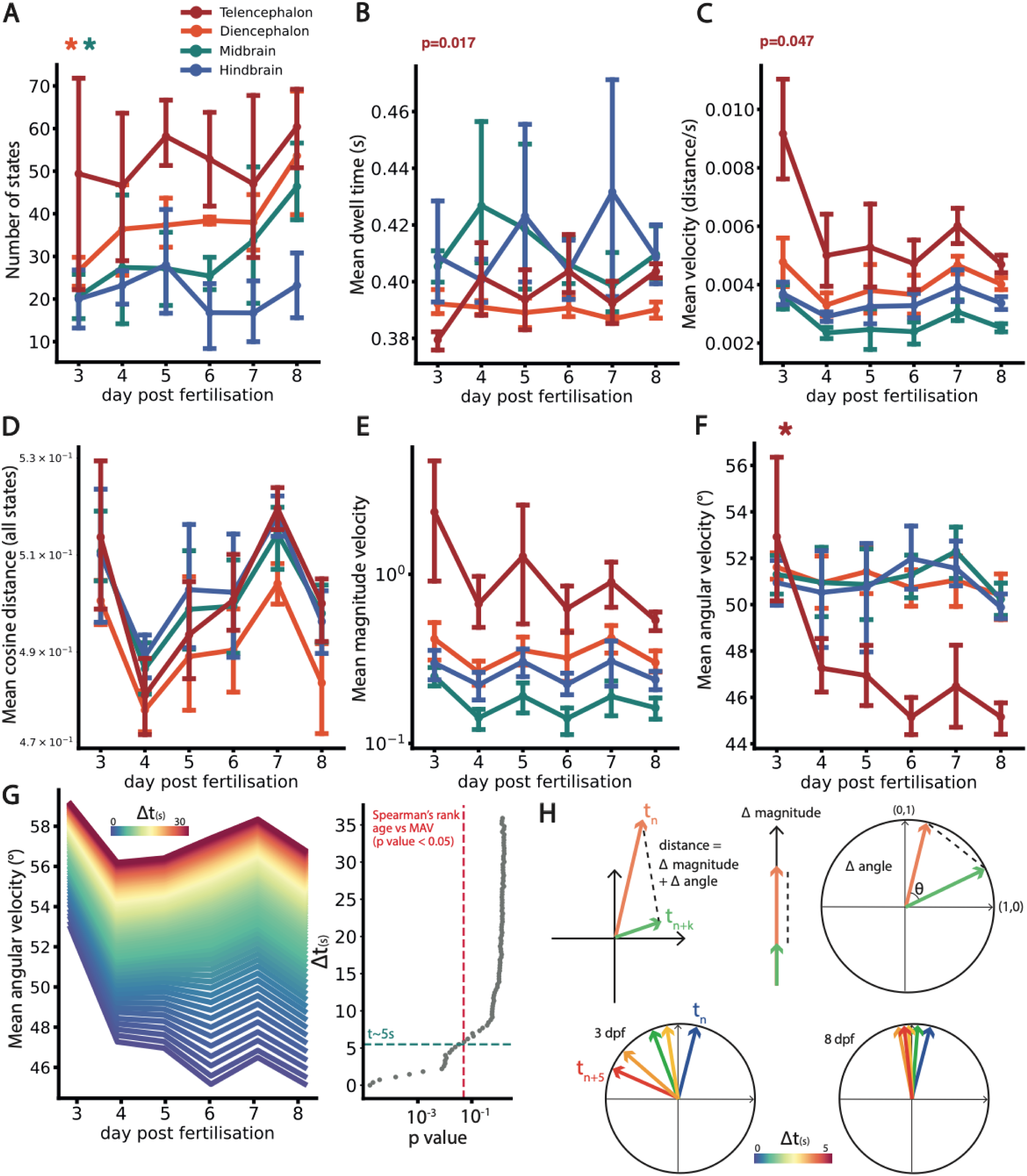
Brain state statistics across development. (A) The number of brain states, (B) the mean dwell time in each state, and (C) the mean velocity (euclidean distance travelled from tn to tn+1), visualised across major brain subdivisions from 3-8 dpf. To decompose the reduction in mean velocity in the telencephalon we measured, (D) the mean cosine distance between all states, (E) the mean magnitude velocity (Δ length of state vectors), and (F) the mean angular velocity (Δ angle between state vectors). (G) The mean angular velocity of the telencephalon plotted as a function of time steps into the future (left), showing a significant age effect until around 5 seconds (right). (H, top) Schematic showing the decomposition of distance travelled between two states into the difference in magnitude and angle of the states. (H, bottom) Schematic illustrating reduced angular velocity at 8dpf compared to 3dpf – the effect is noticeable up to 5 seconds into the future. Data are significant with Bonferroni correction at α = 0.05 for Spearman’s test (*).

To understand what drives longer dwell times and state stability, we calculated the velocity of the dynamics in state space, as a measure of the timescale of the dynamics (D. R. W. Burrows et al., 2023). Velocity was calculated as the mean Euclidean distance between states at t_n_ and t_n+1_ normalised by the number of cells. Here, the telencephalon showed a trend towards a reduction in velocity across development (Spearman’s p=-0.36 p=0.047), while other brain regions showed no clear age effect effects (diencephalon: Spearman’s p=0.06, p=0.74; midbrain: Spearman’s p=-0.20,p=0.28, hindbrain: Spearman’s p=0.11, p=0.57) (**Fig.3C**). Given that the overall effect of such slowing could be masked by multiple population mechanisms we decided to delve further into this slowing effect.

To understand the population mechanisms driving such slower dynamics across development, we decomposed the observed reduction in Euclidean distance travelled into its potential causes: i) An increased similarity of brain states – nearby states reduces the possible distance travelled. ii) An increased similarity of state vector magnitudes – for two state vectors v_n_ and v_n+1_, the difference in vector magnitudes is | ||v_n_|| - ||v_n+1_|| |. This is akin to aligning two vectors and comparing their lengths, which evaluates differences in summed amplitudes across all neurons (**Fig.3H**). iii) An increased similarity in state vector angles – given by the cosine similarity (v_n_. v_n+1_ / ||v_n_|| ||v_n+1_||). This is akin to removing the lengths of the vectors and comparing only their orientation (**Fig.3H**). Firstly, we found no significant age effects when comparing the mean cosine distance across all states (**Fig.3D**) or the vector magnitudes (**Fig.3E**). This indicates that slower dynamics in the telencephalon are not driven by more similar brain states or reduced variance in overall activity. Next, we estimated the mean angular velocity as the mean cosine distance from time *n* to *n*+1 – we found a significant reduction in the angular velocity across development in the telencephalon (Spearman’s p=0.67 p<0.001) from 52.9±3.23 degrees at 3dpf, to 45.2±0.66 degrees at 8dpf. Other brain regions showed no age effects (diencephalon: Spearman’s p=-0.25, p=0.18; midbrain: Spearman’s p=-0.04,p=0.85, hindbrain: Spearman’s p=-0.12 p=0.52) (**Fig.3F**). This demonstrates that the slowing down that we see is caused by more similar vector orientations across sequential state vectors over development (**Fig.3H**). Importantly, in higher dimensions random vectors tend towards orthogonality which would bias cosine distance calculations in networks with different numbers of neurons. To test that observed angular velocity reductions do not occur due to changes in neuron number we re-calculated angular velocity on randomly subsampled data with 300 neurons per fish and brain region, subsampled 50 times per sample. Importantly, as above only the telencephalon shows significant reductions in angular velocity across development (**Supplementary Fig.1**).

This stability in vector orientations across time, which we call rotational stability, occurs due to the emergence of stable relative activity patterns across neurons in the network. Such a rotationally stable regime indicates stronger correlations between sequential brain states across development in the telencephalon. Importantly, this is not driven by the trivial case in which overall activity is stable over time (**Fig.3H**), but instead implies stable linear dependencies across the network in the face of fluctuating activity levels.

We were interested in characterising the timescale of such rotational stability, to understand how far into the future brain states remained rotationally stable, relative to earlier development. To test this, we estimated the mean angular velocity between vectors at time *n* to *n+t*, increasing *t* from 1 to ∼ 30 seconds. When assessing mean angular velocity with age as a function of *t* we find that mean angular velocity decreases across development at small *t,* before plateauing with increasing *t* (**Fig.3G**). Specifically, mean angular velocity shows a significant reduction with age up to ∼ 5.49 seconds into the future (**Fig.3G**). This means that neurons show increased stability in their relative activity patterns across development, visible at a timescale of around 5 seconds. The emergence of such stable dynamics over long timescales indicates the appearance of robust attractor states driven by strongly correlated subnetworks in the telencephalon.

### Rotational Stability Co-occurs with Attractor Stability

Finally, we wanted to validate that the emergence of rotational stability could in fact be driven by stronger attractor states. We reasoned that oscillating signals with converging frequencies and phase alignments would show a tendency to become scaled versions of one another across time. This would mean that sequential state vectors should become more correlated, thus increasing rotational stability. To investigate this we utilized the Kuramoto coupled oscillator model and applied it to our calcium imaging data (Kuramoto, 1975; Penn et al., 2016). We theorised that the coupling strength K, which determines the phase synchronicity among oscillators, would modulate rotational stability. For simplicity, we simulated a network of 100 fully connected oscillators (see Methods, **Fig.4A**). Interestingly, as K approached 0.08, the mean angular velocity of the network exhibited a non-linear decrease before plateauing at approximately 0.13 (**Fig.4B**). Conversely, mean phase coherence increased across this range before plateauing (**Fig.4C**). This suggests that increased coupling strength in oscillating networks is a key driver of both increased phase coherence and rotational stability.

**Figure 4.**
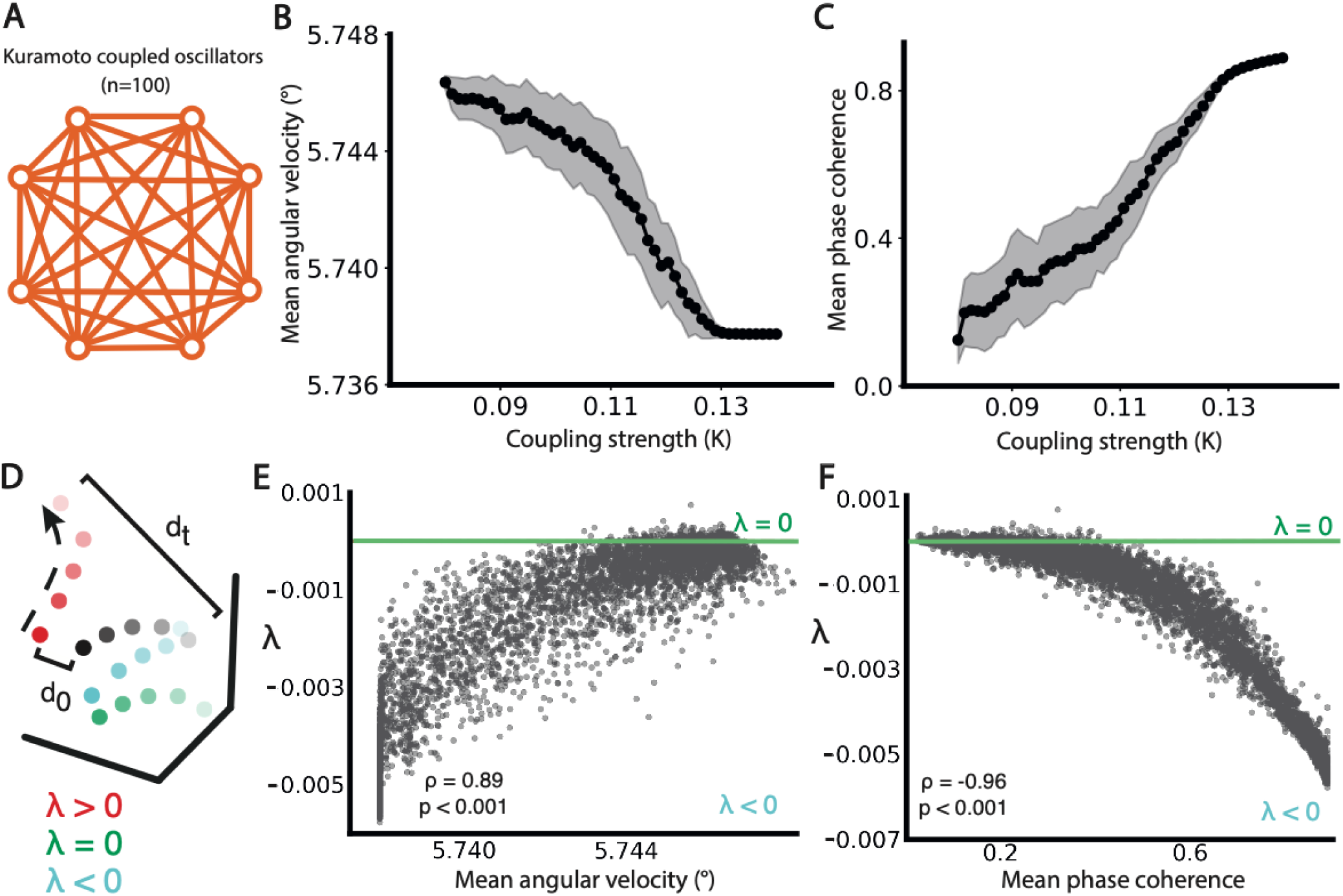
Kuramoto coupled oscillator model of rotational stability. (A) Schematic of Kuramoto oscillator model with 100 fully-connected oscillators with uniform coupling strengths (K). The effect of coupling strength K on mean angular velocity (B), and mean coherence between the oscillators (C). (D) Schematic outlining the meaning of different lyapunov exponent (λ) values. Each colour represents the trajectory over time for a specific initial point in state space, with high to low brightness representing movement in time. λ is the ratio of the difference between two points at the start (d0) and at t (dt). λ > 1: chaotic (red), λ < 1: stable (blue), λ = 1: neutral (green), where λ for each trajectory is calculated against the black trajectory. The relationship between λ and mean angular velocity (E) and mean phase coherence (F). Test statistics and p values are Spearman’s rank correlation test.

To assess stability of the underlying attractor states, we calculated the Lyapunov exponent (*λ)*, a measure of sensitivity to small perturbations in state trajectories (see Methods).

Importantly, if the dynamics are driven by stable attractors then nearby trajectories should get closer over time (*λ* < 0), whereas in more flexible dynamics should result in a conservation of distances across neighbouring trajectories due to weakly attracting dynamics (*λ* ∼ 0) (**Fig.4D**). We found a significant positive correlation between mean angular velocity and *λ* (Spearman’s p=0.89, p<0.001), with angular slowing coinciding with a reduction in *λ*. Interestingly, *λ* remained near 0 before angular velocity reduced, but shifted to negative values as angular velocity decreased (**Fig.4E**). We also observed a strong negative correlation between phase coherence and *λ* (Spearman’s p=−0.96,p<0.001) (**Fig.4F**). This pattern indicates a relationship between rotational stability and dynamical stability – in our model, increasing rotational stability correlated with a divergence away from critical dynamics at a phase transition towards more stable, attracting dynamics.

## Discussion

In this study, we explored the dynamic interplay between stability and flexibility in neuronal populations across the major brain regions at single cell resolution over development. One hypothesis states that the brain progresses towards a phase transition, thus optimising the system for flexibility and sensitivity. Our findings however point towards a more nuanced understanding – in the telencephalon, population dynamics become more scale invariant approaching but not reaching a phase transition, and yet also begin to exhibit stable states over long timescales. These results suggest that the telencephalon evolves towards a regime that balances robust attractor dynamics with the flexibility needed to adapt to changing inputs during critical developmental periods.

Our findings revealed region-specific neuronal excitability changes over early development in larval zebrafish. Only one previous study has examined neuronal firing changes across development in the larval zebrafish, which was limited to the tectum (Avitan et al., 2017). Our findings mirror observed changes in the tectum, with modest increases in excitability around 3-5 dpf. Interestingly, the telencephalon exhibits particularly strong increases in neuronal excitability, indicating a uniquely critical period for telencephalon plasticity in early development. The most dramatic changes occurred from 3 to 4 dpf, which overlaps with periods of axonal and dendritic ramification in telencephalic pallial interneurons (Turner et al., 2022). This period of synaptic remodelling is by no means unique to the telencephalon, with evidence for high synapse turnover in the zebrafish tectum from 3 to 4 dpf as well (Meyer & Smith, 2006). Interestingly, structural data at 6 dpf shows particularly high within-region connectivity in the telencephalon (Kunst et al., 2019). Therefore, such increased neuronal excitability relative to other brain regions might be an emergent property of a regional subnetwork that is becoming more inter-connected across development. This would explain increases in correlations that emerge in the telencephalon. However, little is known about the distinct synaptic constraints on the telencephalon over this period. Future imaging studies are necessary to understand the molecular constraints driving such excitability changes in the telencephalon.

Previous studies have reported that neuronal populations organise their dynamics closer towards a phase transition (criticality) over development (Tetzlaff et al., 2011; Yada et al., 2017). Similarly, our study suggests the expansion of avalanche dynamics in the telencephalon across development, with a higher propensity for larger and longer avalanches that begin to approximate scale invariant distributions. While previous studies used the emergence of scale invariance in avalanche distributions as definitive evidence for criticality, the use of power laws alone is a weak indicator of criticality as such distributions can emerge in a wide array of non-critical systems (Destexhe & Touboul, 2021; Touboul & Destexhe, 2017). In our study we use a variety of features to assess proximity to criticality, namely avalanche power laws, exponent relationships and branching ratios, thus providing a more robust assessment of criticality. In doing so we arrived at a more nuanced understanding of the organisation of population dynamics in development. While avalanches expand in size showing stronger degrees of self-similarity in the telencephalon, branching ratios remain slightly subcritical (∼0.97). This fits with the notion that brain dynamics reside on the slightly subcritical side of the phase transition (D. R. W. Burrows et al., 2023; Priesemann et al., 2014). Given that our study uses single cell recordings with dense coverage across entire brain regions in living systems, while previous studies worked with sparse recordings from cultured neurons, our findings are less likely to be biased by subsampling or neuronal culturing artefacts. These findings provide unique insight into the organisation of neuronal populations towards criticality in early development.

Our population analyses have revealed the functional consequences of emergent population dynamics during early development. Notably, consistent linear dependencies in time lead to rotationally stable dynamics over extended timescales in the telencephalon. This rotational stability demonstrates the emergence of highly correlated sequential brain states that persist for several seconds. Our *in silico* modelling provides evidence that such stability arises from dynamical stability, driven by the formation of more stable attractors that draw the dynamics toward them.

These findings underscore the importance of maintaining stable and robust dynamics throughout development in the telencephalon, while also illustrating a specific population mechanism—rotational stability—through which these stable dynamics can emerge. This raises the question: how do we reconcile the concepts of criticality and rotational stability? Previous theoretical studies have shown that it is possible to balance the stability of subcritical dynamics with the flexibility inherent in criticality. Reverberating regimes, which can occur in the slightly subcritical vicinity of a phase transition (Wilting et al., 2018; Wilting & Priesemann, 2019), exemplify this balance. Operating within this range allows the system to be highly sensitive to changes in synaptic parameters, enabling the network to shift between flexible and stable regimes, whilst also mitigating the risk of entering a supercritical regime, such as during a seizure (D. R. W. Burrows et al., 2023; Liao et al., 2019; Meisel et al., 2012; R. Rosch et al., 2019). Indeed, synaptic plasticity mechanisms can modulate the proximity to a phase transition (Zeraati et al., 2021), such as criticality and more asynchronous or stable dynamics (Li & Shew, 2020). Consequently, developmental programs may be designed to finely tune plasticity to such a subcritical operating range, providing the system with sufficient flexibility to navigate its parameters within a safe range, sufficiently close to stability while remaining close to the phase transition.

## Materials & Methods

### Experimental Models

To capture spontaneous neuronal activity at single cell resolution across the whole brain, we took advantage of the transparency of the larval zebrafish. Transgenic zebrafish larvae expressing a nuclear-localised calcium reporter in all neurons (elavl3:H2B-GCaMP6s) were used to capture neuronal dynamics. To maximise optical transparency, elavl3:H2B-GCaMP6s larvae were out-crossed with melanophore deficient (-/-) roy;nacre mitfa zebrafish (Lister et al., 1999). Zebrafish larvae were raised at 28°C in danieau with a day and night cycle of 12:12 hours, in a petri dish above a gravel environment to mimic visuospatial scenes in the wild (Diana et al., 2019). To capture activity across development, imaging was performed at 3 (n=5), 4 (n=5), 5 (n=6), 6 (n=5), 7 (n=4) and 8 (n=5) days post fertilisation (dpf). This work was approved by the local Animal Care and Use Committee (Kings College London) and was carried out in accordance with the Animals (Experimental Procedures) Act, 1986, under license from the United Kingdom Home Office.

### 2-photon Calcium Imaging

To record single neuron calcium fluctuations across the whole brain, we utilised a fast scanning 2-photon microscope for volumetric imaging. At 6dpf non-anaesthetised larvae were immobilised in agarose and mounted dorsal side up on a raised glass platform, placed in a danieau-filled chamber **(Fig.1A)**. Imaging was performed using a custom built 2-photon microscope (Independent NeuroScience Services), with a Mai Tai HP ultrafast Ti:Sapphire laser (Spectraphysics) at 940nm. Emitted fluorescence was collected by a 16x, 1NA water immersion objective (Nikon) and detected via a gallium arsenide phosphide detector (ThorLabs). Volumetric scanning was performed using a resonance scanner (x-axis) and galvo-mirror (y-axis), with a piezo lens holder (Physik Intrumente) adjusting the z-plane. The objective’s field of view in x and y dimensions covered the entirety of forebrain, thalamic, midbrain and cerebellar regions while covering the rostral portion of the hindbrain.

Volumetric data was collected across 10 focal planes at 15μm intervals in the z dimension, resulting in a frame rate of 2.73 Hz per volume (**Fig.1B**). Images were acquired at a resolution of 512x512 pixels, with a pixel size of 1.05 x 1.03 μm. Imaging was performed for 30 minutes. Larvae were left for 30 minutes in the light to allow the agarose to set and the fish to habituate to its environment.

### Calcium Image Pre-processing

Pre-processing algorithms used to register and segment raw calcium images have been described in detail in previous work (D. R. W. Burrows et al., 2023). Datasets with low baseline fluorescence, drift in the z-plane, or evidence of brain warping were excluded. Overall ∼10,000-20,000 neurons were segmented per fish. Fluorescence traces were then normalised (ΔF/F) and binarized into calcium events using methods described in detail previously (D. R. W. Burrows et al., 2023).

### Regional Labelling of Segmented Cells

To analyse neuronal activity over development across different brain regions, a further registration step was performed to map segmented cells onto their corresponding brain regions. We used the HuC:H2B-RFP image stack from the Zebrafish Brain Browser repository as a reference template (Tabor et al., 2019). Non-linear registration was performed to map a mean image of each 2-photon image plane onto the reference.

Specifically, we used the SyNRA function in ANTSPy to perform Symmetric normalisation: consisting of rigid, affine and deformable transformations (Klein et al., 2009). Custom python code was used to apply the above registration, mapping an image plane to the reference, to its corresponding segmented cell coordinates, to map the cells onto the reference. Each cell was labelled according to its x-y-z position in the reference space, according to the Gupta atlas which consists of genetic labels from 210 transgenic lines (Gupta et al., 2018). This enabled the assignment of each segmented neuron to one of the four major brain sub-divisions: the telencephalon, diencephalon, midbrain and hindbrain (**Fig.1C**).

### Dimensionality Estimation

In order to quantify the covariance across the entire population, we used a measure of dimensionality developed previously (Recanatesi et al., 2019), defined as

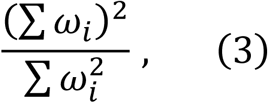

where *ωi* captures the eigenvalue of the *i^th^* principal component. Here, the dimensionality captures the extent to which the variance over the principal components is shared equally (high dimensionality) or dominated by a few components (low dimensionality). If one large, highly correlated subnetwork dominates the dynamics then the variance of the data could be captured with only a few eigenvectors, while most remaining orthogonal eigenvectors would be redundant resulting in a low value in (3). In this case, high population covariance would lead to a low dimensionality. To calculate eigenvalues for (3), PCA was performed on a covariance matrix across all neurons for a given brain region and animal.

### Avalanche Estimation

To estimate the propagation of activity through the network, we use the neuronal avalanche framework which describes how local bursts of activity spread through the system (Beggs & Plenz, 2003). To estimate neuronal avalanche dynamics we need to infer the spread of activity, without knowing the underlying synaptic connectivity of the network. To do this we used an algorithm previously developed by our lab for estimating avalanche dynamics in single cell, whole brain recordings (D. R. W. Burrows et al., 2023). For detailed methodology and parameter settings please refer to the previous work.

First, we assume that neuron *u* can only activate other neurons in the population *P* that lie within a neighborhood *N_u,v_*where *v* is the set of the closest *k*% of cells to *u*. This introduces a spatial constraint to avalanche propagation such that activity can only flow between putatively connected neurons, preventing disparate cascades combining into one large avalanche. An avalanche begins when at least *n* cells within a neighborhood *N_u,v_* are active at *t_x_*, that were not part of an avalanche in the previous timestep. Here, we label the neurons currently active at *t_x_*which belong to the avalanche as the set *A_x_* = {*a,b,c*…*z*}. All active neurons of *P* at *tx* connected to any of *Ax* via a neighborhood are included into the set *Ax*.

At each subsequent step avalanche propagation iterates as follows:

1. If at least one neuron that was part of avalanche *A_x_* is also active at *t_x+1_* then the avalanche continues in time, forming the set *A_x+1_* = {*a,b,c*…*z*}.
2. Any neurons from *P* active at *t_x+1_* that are connected to any of *A_x+1_* via a neighbourhood are included into the set *A_x+1_*.

Once step 1 is no longer satisfied the avalanche terminates. Avalanches whose active neighbourhoods converge are grouped into a single avalanche. Avalanche size was calculated as the total number of calcium events during the avalanche. Avalanche duration was the number of time steps for which the avalanche was active.

### Power Law Estimation

A key property of systems at a phase transition (criticality) is that avalanche dynamics become scale invariant (Perković et al., 1995; Sethna et al., 2001). These scale invariant avalanches give rise to power law distributions of avalanche size *S* and duration *T* with the following form:

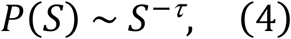

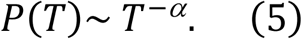

To test for the presence of power law distributions we used an importance sampling approach applied in previous work (D. R. W. Burrows et al., 2023). Here we compared the likelihood of our data being generated by a power-law compared with a log normal distribution, an alternative heavy tailed distribution conventionally used as an alternative hypothesis (Clauset et al., 2009). The log likelihood ratio was calculated as the maximum likelihood of a power law minus the maximum likelihood of a lognormal distribution, with positive values indicating more evidence for a power law distribution. Avalanche power law exponents were estimated as the maximum likelihood of λ given the data.

### Exponent Relationships

While power laws are indeed found in avalanches at criticality, they can emerge in non-critical and stochastic systems (Miller, 1957). Therefore, other markers are needed as further proof of critical dynamics. Another key property of criticality, is that if equations (2) and (3) hold, then there must be a scaling relationship between avalanche size *S* and duration *T*, as

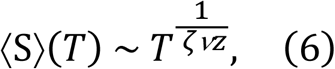

where〈S〉(T) is the average size for a given duration, and *1/ζνz* is a combination of critical exponents, where *ζ* defines the dependence of the power law cut off, *ν* defines the divergence of the size of avalanches, and *νz* defines the divergence of the duration of avalanches, as the critical point is approached (Perković et al., 1995). Put simply, the expected size of an avalanche should scale with its duration as a power law, with exponent 1/*ζ*νz. This occurs because many critical systems exhibit self-similarity across space and time scales, such that a single scaling exponent can describe the relationship between size *S* and *T* at different scales. For simplicity we refer to 1/*ζ*νz as γ. Importantly, at criticality the relationship between the scaling exponent γ and avalanche exponents τ and α should follow:

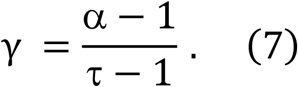

This equality or exponent relation, can be reached by rescaling the function 〈S〉(T), and occurs when a scaling relationship exists between avalanche size and duration as expected at criticality (Burrows, 2022; Sethna et al., 2001; Touboul & Destexhe, 2017). To assess the presence of this critical exponent relation, we make use of the Deviation from Criticality Coefficient (DCC) (Ma et al., 2019) to assess the divergence of observed scaling exponents from the form given in equation 5. DCC is defined as the absolute difference between the scaling exponent γ, measured from the slope of〈S〉(T), and the fraction (α - 1)/(τ - 1).

### Kuramoto Coupled Oscillator Model

To model the relationship between coupling strength, angular velocity and dynamical stability we constructed a Kuramoto coupled oscillator model of 100 fully connected units (Kuramoto, 1975). The dynamics are defined by:

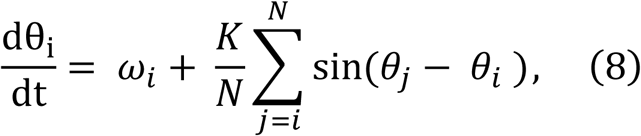

where *θi* is the phase of the ith oscillator, *ωi* is the intrinsic frequency of the ith oscillator, *K* is the coupling strength between all oscillators, *N* is the number of oscillators and *sin(θj - θi)* is the synchronicity term that attracts oscillators towards the same phase. We randomly sampled intrinsic frequencies *ω* for each oscillator from a uniform distribution with range 0.9 to 1.1. Initial phases for each oscillator were randomly sampled from a uniform distribution with range 0 to 2π. Angular velocity was estimated by first calculating the sin and cosine of the phases of the oscillators, which were concatenated into a vector of 200 dimensions, and then computing the cosine distance across states. To study angular velocity we tuned the model to the point where altering K led to a change in mean angular velocity (K∼0.08). We linearly sampled 50 K values between 0.08 to 0.14, with 100 iterations at each K value for simulations. Models were simulated for 200 time steps before estimating mean angular velocity across consecutive states.

To estimate the Lyapunov exponent (λ) we added a small perturbation to the network at t =200. λ was estimated as

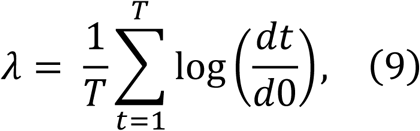

where *d0* and *dt* are the distances between unperturbed and perturbed networks at time 0 and *t*, respectively. *T* was estimated 100 time steps into the future.

### Statistical Tests

D’agostino’s K2 test was used to test for normality in data distributions (α = 0.05). Independent t-tests or Mann Whitney U tests were used to compare firing statistics in the telencephalon with other brain regions, in cases of normality and non-normality respectively (α = 0.05). Spearman’s rank correlation test was used to identify significant relationships between neuronal properties and age (α = 0.05). For testing different brain regions, Bonferroni correction was applied at a threshold.

### Software

Data was analysed using custom code written in Python. Image registration and cell segmentation was performed using Suite2p (Pachitariu et al., 2017). Nonrigid registration was performed using ANTSPy (Klein et al., 2009). Hidden Markov Models were fit using software from Diana et al., 2019. Avalanches were estimated using software from (D. R. W. Burrows et al., 2023). Branching parameters were estimated using MR. Estimator (Spitzner et al., 2021). Statistical hypothesis tests were performed using scipy. Graphs were generated using matplotlib and seaborn.

### Resources

Information and requests for resources should be directed to the corresponding author. Full datasets will be made accessible after publication.

Custom written Python code can be accessed at: https://github.com/dmnburrows/multiscale_dev_dynamics https://github.com/dmnburrows/criticality https://github.com/dmnburrows/img_process

## Acknowledgements & Funding Sources

D.R.W.B. was supported by a Medical Research Council-Sackler PhD Fellowship. R.E.R, received funding from Wellcome (209164/Z/17/Z).

---

**Supplementary Figure 1.**
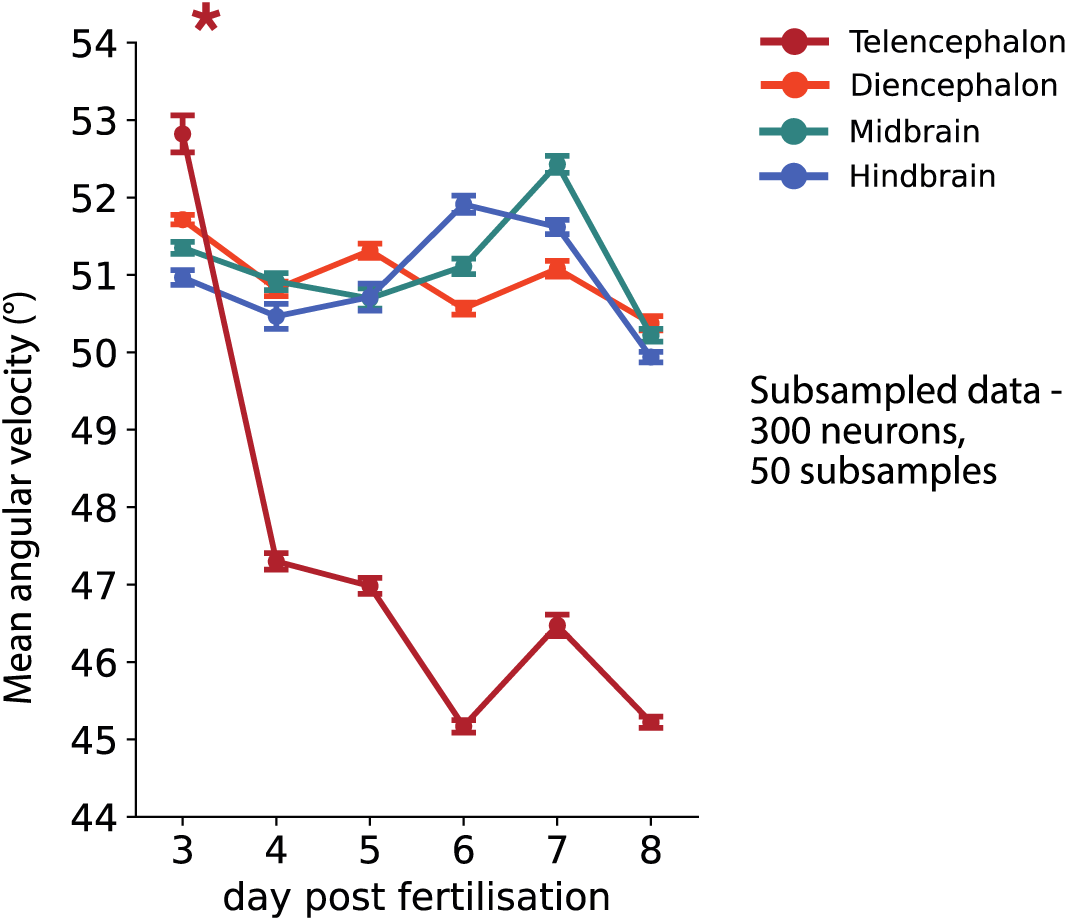
Mean angular velocity in subsampled data. Data was subsampled to 300 neurons, randomly sampled 50 times per sample, before calculating mean angular velocity. Data are significant with Bonferroni correction at α = 0.05 for Spearman’s test (*).

